# Wall teichoic acid glycosylation shapes surface and secreted protein distribution in *Listeria monocytogenes*

**DOI:** 10.64898/2026.03.30.715212

**Authors:** Gonçalo Matos, Ricardo Monteiro, Didier Cabanes

**Author notes:** Corresponding authors: Ricardo Monteiro and Didier Cabanes, i3S (Instituto de Investigação e Inovação em Saúde), Rua Alfredo Allen 208, 4200-135 Porto, Portugal., and.

## Abstract

*Listeria monocytogenes* relies on a tightly controlled set of surface-associated and secreted proteins to mediate host interaction and infection. The correct localization and exposure of these proteins at the bacterial surface are critical for virulence, yet the role of cell wall components in organizing this process remains incompletely understood. In particular, wall teichoic acid (WTA) glycosylation has been implicated in anchoring and function of selected surface proteins, but its global impact on protein distribution across the bacterial cell envelope is unclear.

Here, we performed a comprehensive proteomic analysis to investigate how WTA glycosylation influences protein distribution in *L. monocytogenes*. Using isogenic mutants lacking rhamnose (Δ*rmlT*) or GlcNAc (Δ*lmo1079*) WTA glycosylation, we compared the exoproteome, the surface-accessible proteome and the surface-exposed proteome.

Loss of WTA glycosylation did not result in a global disruption of the surface proteome but instead induced a redistribution of proteins across extracellular and surface-associated fractions. This effect was dependent on protein anchoring mechanisms, with limited changes observed for LPXTG-anchored proteins, moderate effects on non-covalently associated proteins, and a marked enrichment of lipoproteins in the surface-exposed proteome, particularly in the Δ*lmo1079* mutant. In parallel, virulence-associated proteins displayed altered accessibility and exposure, with a progressive shift towards increased surface localization and a combination of shared and mutant-specific responses. This global effect was supported by functional annotation, which revealed that the affected proteins were associated with similar biological processes across fractions, highlighting a broad rather than pathway-specific impact of WTA glycosylation loss

Together, these findings indicate that WTA glycosylation plays a key role in organizing the bacterial surface by modulating protein retention, exposure and release. Rather than affecting specific proteins, WTA glycosylation broadly shapes the spatial distribution of proteins across the cell envelope, with potential consequences for host– pathogen interactions.

## Introduction

*Listeria monocytogenes* is a Gram-positive foodborne pathogen responsible for severe invasive infections, particularly in immunocompromised individuals, the elderly, and pregnant women^1^. Its ability to successfully colonize and invade host cells relies on a tightly controlled set of surface-associated and secreted proteins that mediate adhesion, invasion, and intracellular survival^2^. The proper localization and exposure of these proteins at the bacterial surface are therefore critical determinants of virulence and host–pathogen interactions^3^. In this context, the bacterial surface represents a highly dynamic interface where protein positioning directly influences functional activity and interaction with the host environment^4^.

In Gram-positive bacteria, the cell envelope is a complex and dynamic structure composed of peptidoglycan and associated polymers, including WTAs. These anionic glycopolymers are covalently linked to the peptidoglycan and extend through the cell wall, contributing not only to structural integrity but also to the physicochemical properties of the bacterial surface^5,6^. Importantly, the Gram-positive cell wall constitutes a dense and highly organized matrix that can restrict or allow the accessibility of proteins and external molecules, thereby shaping the functional exposure of the bacterial surface^7^. In *L. monocytogenes*, WTAs are further modified by specific glycosyltransferases, leading to the addition of sugar residues such as rhamnose and GlcNAc^8^. These glycosylations have been implicated in many processes, including virulence, resistance to antimicrobial compounds, and interactions with the host environment^9,10^.

Beyond their structural role, increasing evidence suggests that WTA glycosylation contributes to the spatial organization of surface-associated proteins also in other pathogenic and non-pathogenic bacteria, such as *Staphyloccous aureus*^11,12^ and *B. subtillis*^*13*^. Surface proteins in Gram-positive bacteria are anchored through diverse mechanisms, including covalent attachment via sortase-dependent LPXTG motifs, non-covalent interactions with cell wall polymers such as WTAs, and membrane anchoring as lipoproteins, which may be differentially affected by changes in cell wall composition^14^. In particular, non-covalently associated proteins, including key virulence factors such as internalins and autolysins, rely on interactions with WTAs for their proper anchoring at the bacterial surface^15^. Alterations in WTA composition can therefore affect the localization, accessibility, and release of these proteins, with potential consequences for bacterial physiology and pathogenicity^16^. Despite these insights, most studies to date have focused on individual proteins or specific mechanisms^17–19^, and a comprehensive understanding of how WTA glycosylation globally influences protein distribution across the bacterial cell wall remains limited. Dissecting protein localization across the bacterial surface therefore requires complementary experimental approaches capable of distinguishing between released proteins, surface-accessible proteins, and those directly exposed at the cell envelope surface^20^. In this study, we aimed to systematically investigate the impact of WTA glycosylation on protein distribution across extracellular and surface-associated compartments in *L. monocytogenes*. Using isogenic mutants lacking the glycosyltransferases responsible for rhamnose (*ΔrmlT*) and GlcNAc (*Δlmo1079*) modifications, we performed a comparative proteomic analysis of the exoproteome, the surface-accessible proteome and the surface-exposed proteome, providing a framework for better understanding how WTA glycosylation contributes to the organization of the bacterial cell envelope.

## Results

### Profiling of *Listeria monocytogenes* surface and secreted proteins

The present study aims to characterize the impact of WTA glycosylation on the surface-exposed and secreted protein landscape of *L. monocytogenes* EGDe. A comprehensive proteomic analysis was performed to define the protein of three *Listeria monocytogenes* strains, namely, the wild-type EGDe and its two isogenic knock-out mutants of *ΔrmlT* and *Δlmo1079*, lacking the glycosyltransferases for rhamnose and GlcNAc respectively in three experimental conditions: i) exoproteome, ii) surface-exposed proteome (shaving) and surface-accessible proteome (biotinylation) (**Figure 1**).

**Figure 1:**
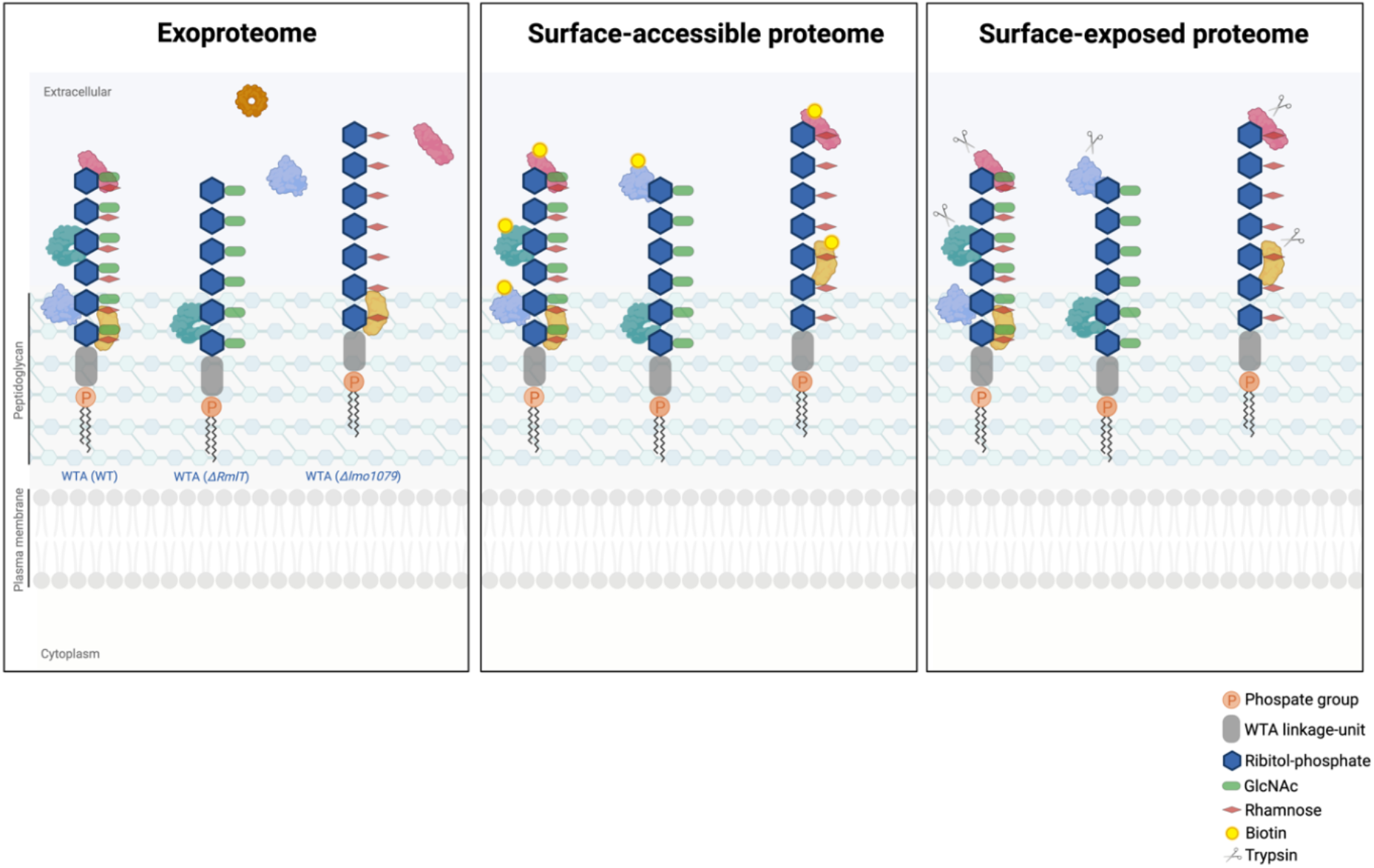
Overview of the experimental strategies used to define distinct proteome fractions. The exoproteome, surface-accessible proteome, and surface-exposed proteome are depicted, illustrating the underlying principles of each method. The exoproteome comprises proteins released into the extracellular milieu, including secreted proteins and those that may fail to remain cell wall–associated. Surface-accessible proteins are identified by biotinylation of exposed residues, whereas surface-exposed proteins are defined by protease accessibility (trypsin shaving). The cell wall is represented with strain-specific WTA glycosylation patterns, including rhamnose and GlcNAc modifications in WT, absence of rhamnose in *ΔrmlT*, and absence of GlcNAc in *Δlmo1079*. Changes in cell wall architecture, such as altered glycosylation, may influence protein anchoring and accessibility. Proteins are represented schematically and do not reflect their actual distribution or abundance.

Across all experimental conditions, a total of 513 proteins were identified in the exoproteome, 1547 in the surface-accessible proteome, and 1146 in the surface-exposed proteome (**Table S1**). Comparable numbers of proteins were detected for the WT strain and the Δ*rmlT* and Δ*lmo1079* mutants in each fraction, indicating consistent proteome coverage across strains. High reproducibility was observed between biological replicates (three), with consistent protein identification across independent experiments within each fraction. Only proteins detected with FDR ≤ 0.01 and >2 peptides were retained for subsequent analyses.

To assess the global distribution and reproducibility of the identified protein populations, Venn diagrams were generated comparing proteins detected in the exoproteome, the surface-accessible proteome, and the surface-exposed proteome of the three different *Listeria* strains (**Figure 2A**). In all cases, the great majority of proteins (>90 %) were shared across conditions, indicating a high degree of consistency in protein identification. This extensive overlap suggests that the effects observed in subsequent analyses primarily reflect quantitative differences in protein abundance rather than stochastic detection or technical variability. At the same time, each approach revealed a smaller subset of fraction-specific proteins, consistent with differences in protein release, accessibility, and exposure captured by the three complementary methods.

**Figure 2:**
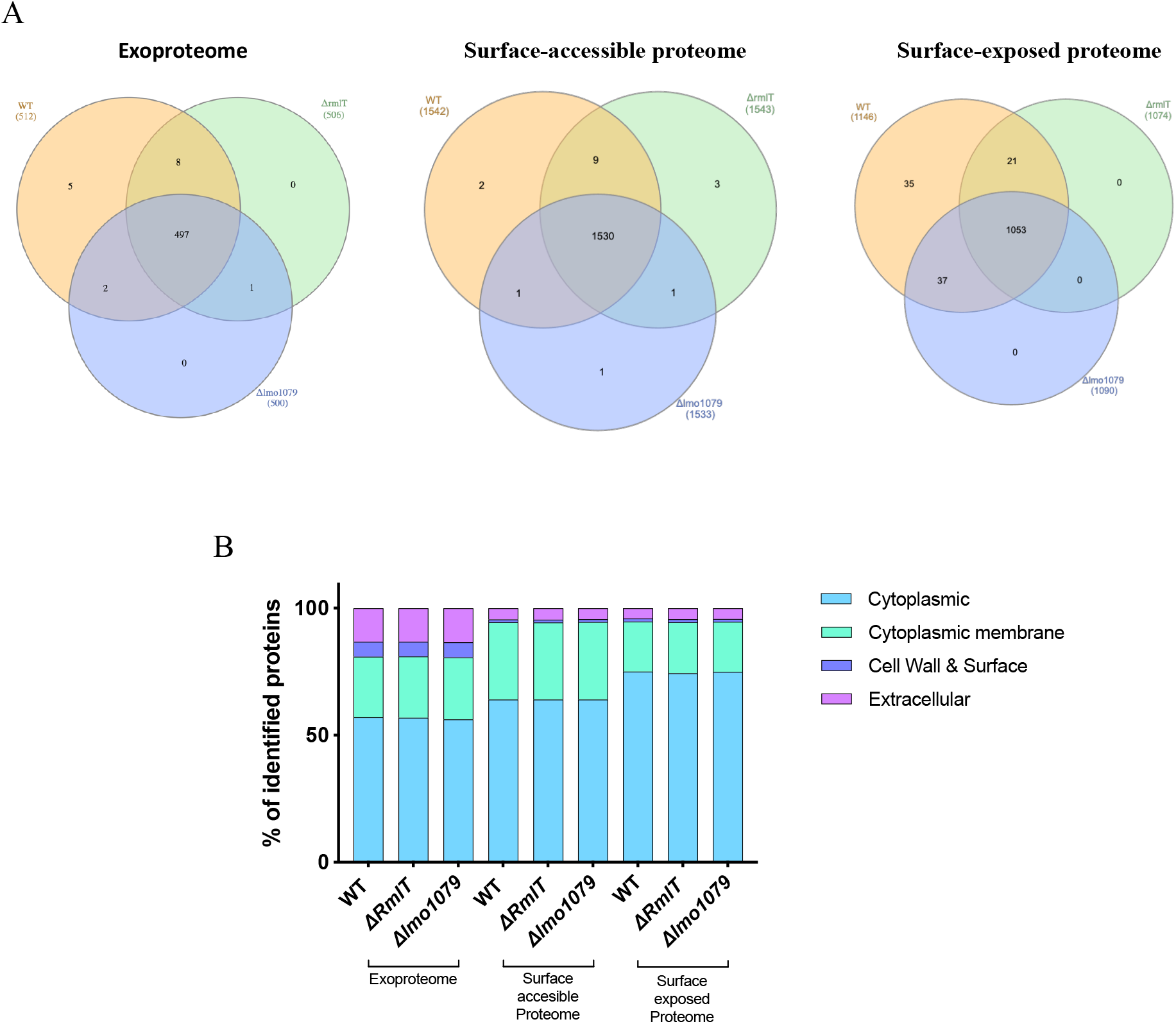
Global overview of protein distribution and predicted localization across extracellular and surface-associated proteomes of *Listeria monocytogenes*. (A) Venn diagrams showing the overlap and fraction-specific distribution of proteins identified in the exoproteome (culture supernatant), the surface-accessible proteome obtained by chemical biotinylation, and the surface-exposed proteome obtained by protease shaving of intact cells. The diagrams highlight the extensive overlap between datasets as well as proteins uniquely detected by each experimental approach. (B) Distribution of identified proteins according to predicted cellular localization and secretion pathway in the exoproteome, surface-accessible proteome, and surface-exposed proteome. Proteins were classified as cytoplasmic, membrane-associated, secreted via canonical pathways, or moonlighting proteins based on bioinformatic annotation. The relative proportions are shown for the WT strain and the Δ*rmlT* and Δ*lmo1079* mutants.

Following the global comparison of protein overlap between fractions, the identified datasets were further characterized according to predicted cellular localization (**Figure 2B**). In the exoproteome, proteins annotated as cytoplasmic constituted the largest proportion of identified proteins, representing 57% in the WT strain, 56.9% in the Δ*rmlT* mutant, and 56.2% in the Δ*lmo1079* mutant, while membrane-associated proteins accounted for 23.8– 24.4% in all strains. Proteins predicted to follow canonical secretion pathways (cell wall and surface) represented a smaller fraction of the exoproteome, ranging from 5.7% to 6.0%. In the surface-accessible proteome, cytoplasmic proteins accounted for approximately 64% of the identified proteins, followed by membrane-associated proteins (30.3–30.5%) and a minor proportion of cell wall associated proteins (approximately 1.1%). In the surface-exposed proteome, cytoplasmic proteins represented the majority of identified proteins (74–75%), with membrane-associated proteins accounting for around 20% and cell wall associated proteins for approximately 1%. In all experimental conditions, comparable distributions were observed across the WT strain and both glycosylation mutants, indicating that differences observed in subsequent analyses primarily reflect changes in protein abundance rather than major shifts in proteome composition.

Together, these results establish the overall composition and consistency of the exoproteome and surface-associated proteomes across the three strains. Given the comparable protein distributions observed between the WT and glycosylation mutants, subsequent analyses focused on quantitative differences in protein abundance to assess how WTA glycosylation affects protein anchoring to the cell wall and modulates the localization, surface exposure, secretion, and release of cell wall-associated proteins, particularly virulence factors

### *L. monocytogenes* alters its subproteomes in response to WTA glycosylation loss

To determine the specific impact of WTA glycosylation on surface and extracellular protein dynamic, we next performed a quantitative comparison of the exoproteome, surface-accessible proteome, and surface-exposed proteome between the WT strain and each glycosylation mutant. Differential abundance analyses were conducted independently for Δ*rmlT* and Δ*lmo1079* in order to define the protein subsets affected by the loss of rhamnose or GlcNAc decoration, respectively. These analyses allowed the identification of proteins significantly increased or decreased in each mutant relative to the WT (**Table S2**).

Differentially abundant proteins were defined using a threshold of 2 (log2 abundance ratio) and an adjusted p-value ≤ 0.01. Based on these criteria, 13 proteins were significantly altered in the exoproteome of the Δ*rmlT* mutant, whereas 16 were altered in Δ*lmo1079*. In the surface-accessible proteome, 47 and 66 proteins were differentially abundant in Δ*rmlT* and Δ*lmo1079*, respectively. Similarly, in the surface-exposed proteome obtained by shaving, 113 proteins were significantly changed in Δ*rmlT* and 126 in Δ*lmo1079* compared to the WT strain.

In the exoproteome, the Δ*rmlT* mutant exhibited six proteins with decreased abundance and seven with increased abundance relative to WT, whereas the Δ*lmo1079* mutant showed a predominance of decreased proteins (13 down and three up). In the surface-accessible proteome, 22 proteins were decreased and 25 increased in Δ*rmlT*, while Δ*lmo1079* displayed 33 decreased and 33 increased proteins in abundance. The most pronounced alterations were observed in the surface-exposed proteome. In this fraction, 87 proteins were decreased and 26 increased in Δ*rmlT*, whereas 65 proteins were decreased and 61 increased in Δ*lmo1079*. These results indicate that the impact of WTA glycosylation loss is most extensive at the level of the surface-exposed proteome, with a marked enrichment of decreased proteins, particularly in the Δ*rmlT* mutant.

To determine whether the observed alterations were mutant-specific or reflected a common response to WTA glycosylation loss, the sets of differentially abundant proteins identified in Δ*rmlT* and Δ*lmo1079* were compared for each fraction. In the exoproteome, seven proteins were commonly altered in both mutants. In the surface-accessible proteome, 22 proteins were shared between Δ*rmlT* and Δ*lmo1079*. The strongest overlap was observed in the surface-exposed proteome, where 57 proteins were consistently altered in both mutants. Notably, the majority of these shared proteins displayed reduced abundance in both mutants, indicating a common dependence on WTA glycosylation for surface exposure.

To gain insight into the biological processes and molecular functions associated with WTA glycosylation– dependent alterations, Gene Ontology (GO) annotation analysis of biological processes (**Figure 3A**) and molecular functions (**Figure 3B**) was performed on the sets of differentially abundant proteins identified in each fraction and strain. Although no GO term reached statistical significance after multiple testing correction, functional categorization revealed consistent biological trends across fractions.

**Figure 3.**
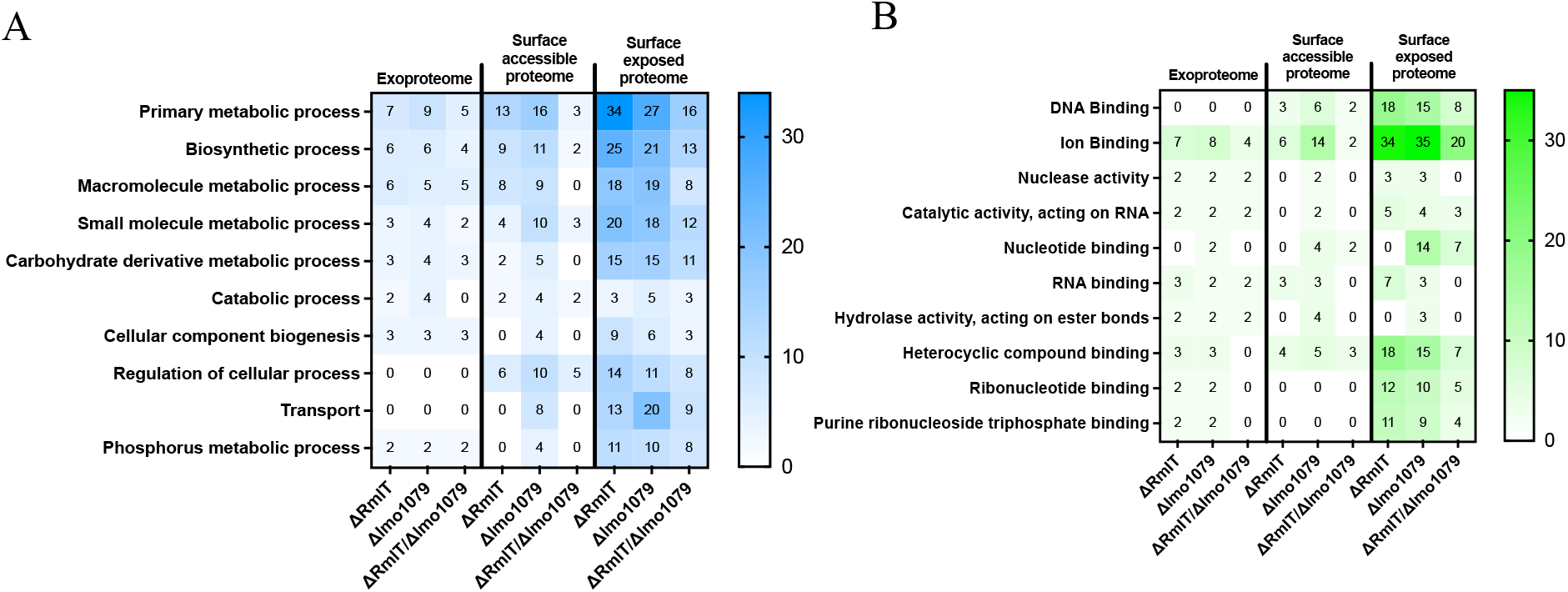
Functional categorization of proteins altered in *ΔrmlT* and *Δlmo1079*, including proteins commonly altered in both mutants, across extracellular and surface-associated fractions. (A) Distribution of Gene Ontology (GO) biological processes associated with proteins altered in the extracellular, surface-accessible, and surface-exposed fractions. (B) Distribution of GO molecular functions for the same protein sets. GO terms were ranked according to protein counts, and only the top 10 most represented terms were retained for visualization. Number indicate the protein’s count assigned to each GO category.

In the exoproteome, proteins altered in *ΔrmlT* were predominantly associated with primary metabolic processes (7 proteins), followed by biosynthetic processes and macromolecule metabolic processes (6 proteins each). At the molecular function level, ion binding was the most frequent category (seven proteins), together with RNA binding and heterocyclic compound binding (3 proteins each). In *Δlmo1079*, a comparable functional distribution was observed, with altered proteins mainly associated with primary metabolic processes (9 proteins), followed by biosynthetic processes (6 proteins) and macromolecule metabolic processes (5 proteins). At the molecular function level, ion binding again represented the most prevalent activity (8 proteins). Among the proteins commonly altered in both mutants, the same biological processes predominated, and ion binding remained the most represented molecular function.

A similar distribution was observed in the surface-accessible proteome. In *ΔrmlT*, altered proteins were mainly associated with primary metabolic processes (13 proteins), followed by biosynthetic processes (9 proteins) and macromolecule metabolic processes (8 proteins). Molecular function annotations were again dominated by ion binding (6 proteins), together with heterocyclic compound binding (4 proteins) and DNA- and RNA-binding (3 proteins each). In Δ*lmo1079*, primary metabolic processes also represented the largest category (16 proteins), followed by biosynthetic processes (11 proteins) and small molecule metabolic processes (10 proteins). Notably, proteins associated with regulation of cellular processes (10 proteins) and transport-related functions (8 proteins) were also annotated. At the molecular function level, ion binding again represented the most frequent category (14 proteins), together with heterocyclic compound binding and DNA binding. Interestingly, among the proteins commonly altered in both mutants in this fraction, regulation of cellular processes was the predominant biological category (5 proteins), while heterocyclic compound binding represented the most frequent molecular function. The strongest functional redistribution was observed in the surface-exposed proteome. In *ΔrmlT*, altered proteins were mainly associated with primary metabolic processes (34 proteins), followed by biosynthetic processes (25 proteins) and macromolecule metabolic processes (18 proteins). Molecular function analysis again highlighted ion binding as the predominant category (34 proteins), together with DNA binding and heterocyclic compound binding (18 proteins each). In *Δlmo1079*, the same biological categories were predominant, with primary metabolic processes representing the main affected category (27 proteins), although transport-related processes were also well represented (20 proteins). At the molecular function level, ion binding again represented the largest category (35 proteins), together with DNA binding and heterocyclic compound binding. Among the proteins commonly altered in both mutants in the surface accessible fraction, primary metabolic and biosynthetic processes were again predominant, and a substantial proportion displayed ion-binding activity.

Across all fractions, altered proteins were predominantly associated with primary metabolic and biosynthetic processes and frequently displayed ion-binding activities. This recurrent functional enrichment suggests that WTA glycosylation loss broadly affects the abundance and surface localization of enzymes and regulatory proteins linked to central cellular metabolism.

### WTA glycosylation loss reshapes the surface and secreted proteomes

To investigate whether the proteins altered upon WTA glycosylation loss belonged to specific classes of cell surface–associated proteins, the differentially abundant proteins identified in each fraction were categorized according to their predicted anchoring mechanisms (**Figure 4**).

**Figure 4.**
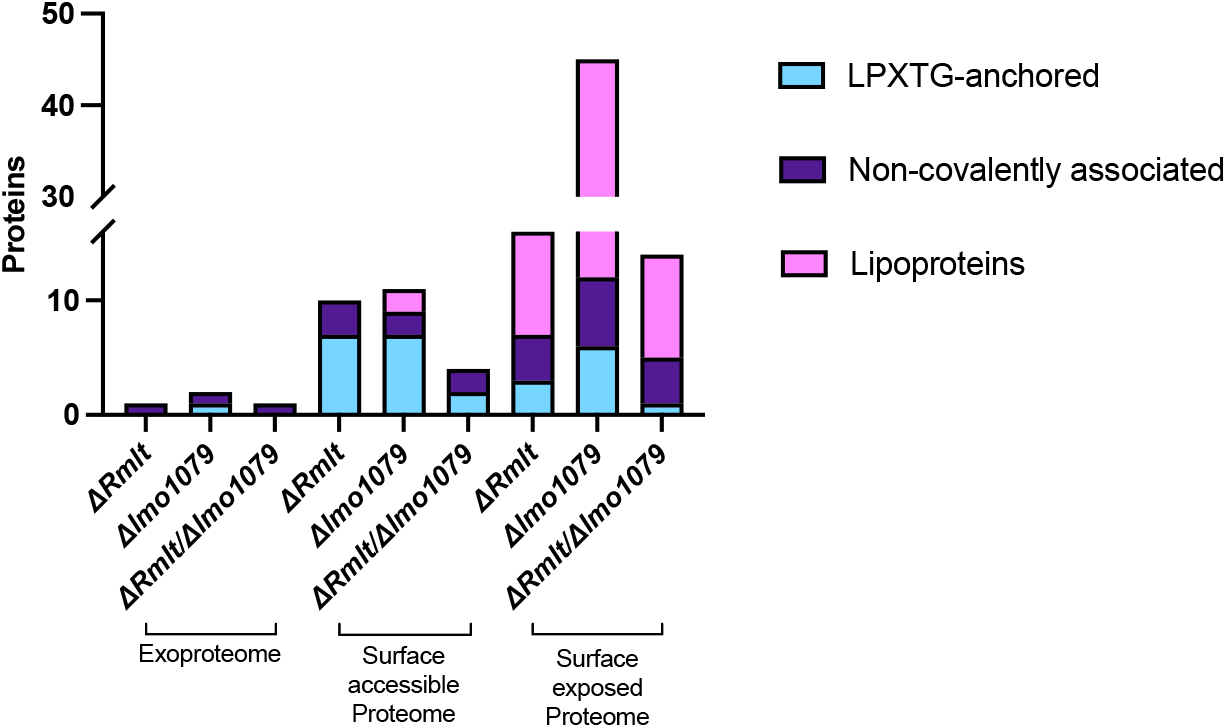
WTA glycosylation loss differentially affects protein anchoring classes and virulence-associated proteins across extracellular and surface-associated fractions. Distribution of differentially abundant proteins according to predicted anchoring mechanism in the extracellular, surface-accessible, and surface-exposed fractions of *ΔrmlT* and *Δlmo1079* mutants relative to WT. Proteins were classified as LPXTG-anchored, non-covalently cell wall–associated, or lipoproteins.

Classification was based on bioinformatic predictions of secretion signals, transmembrane regions, and canonical anchoring motifs, allowing proteins to be grouped as LPXTG-anchored proteins, non-covalently cell wall– associated proteins or lipoproteins (**Table 1**).

**Table 1.**
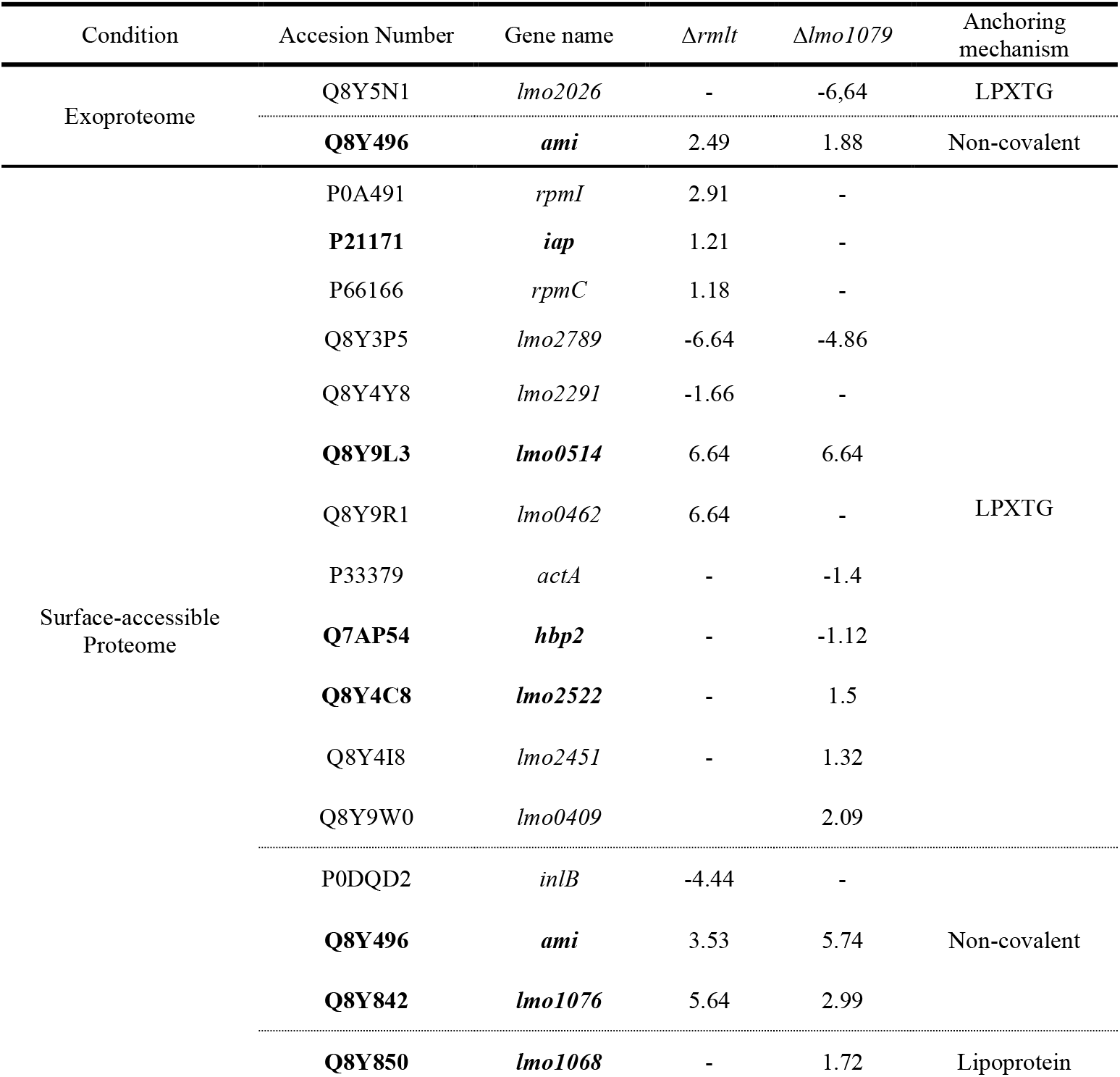
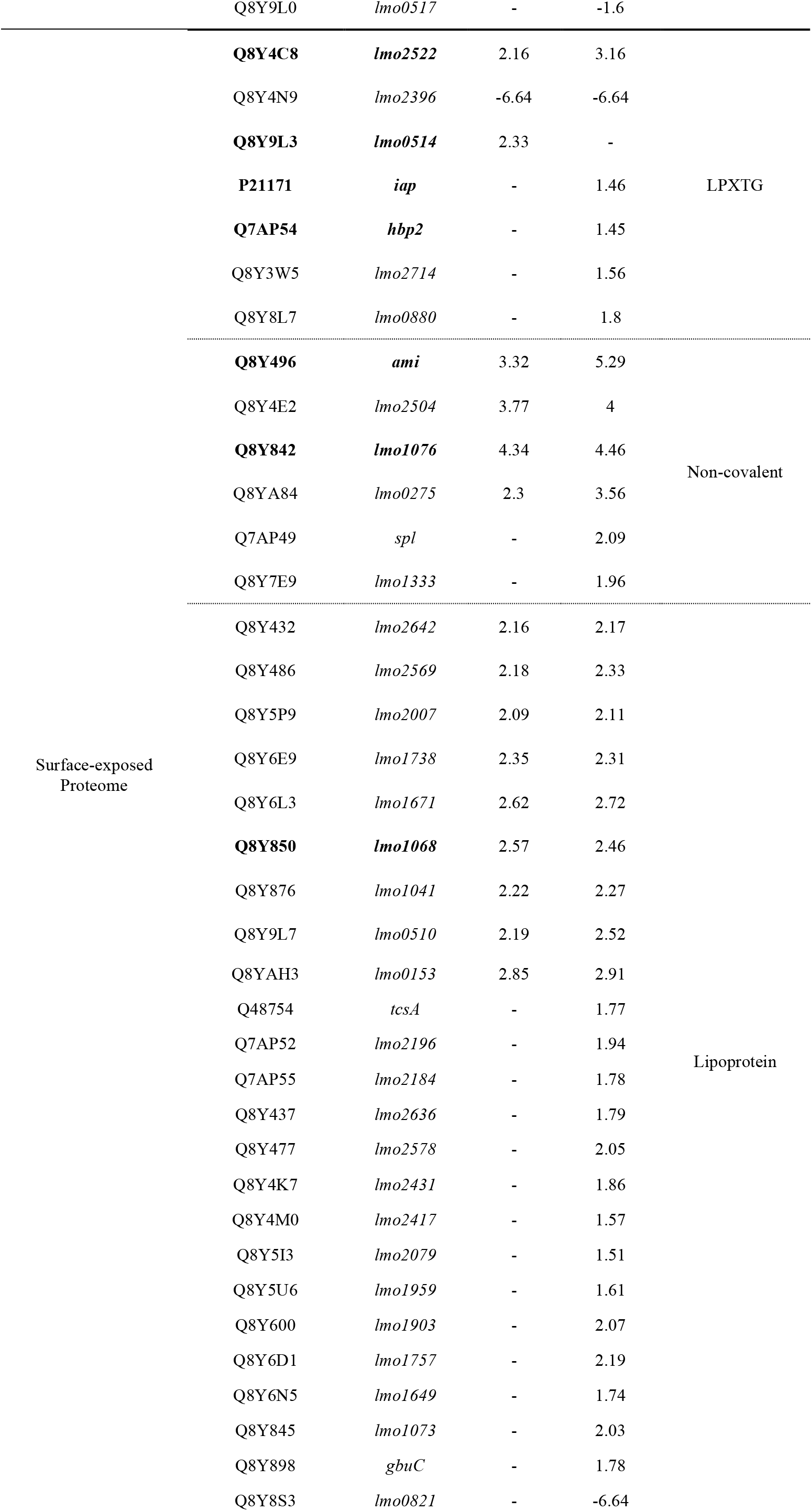
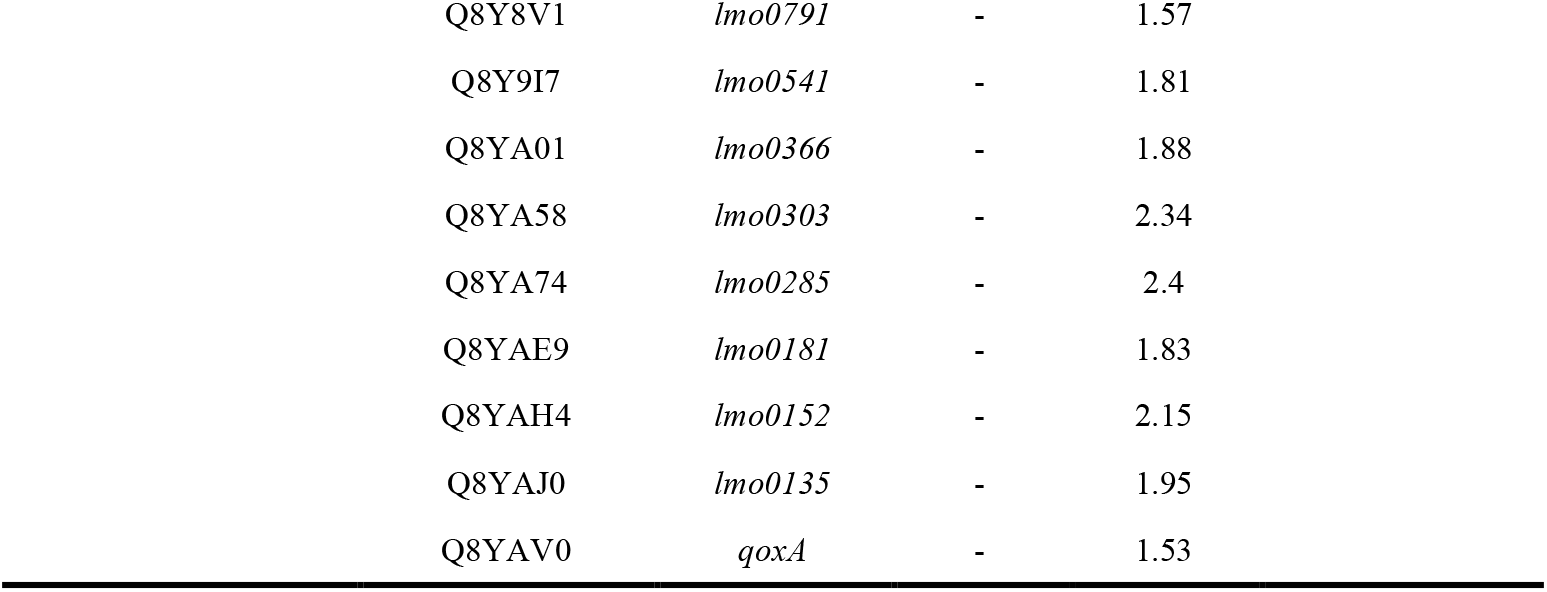
Proteins classification regarding the cell wall anchoring class. Proteins were classified based on in silico predictions of signal peptides, transmembrane helices, and canonical cell wall–anchoring motifs. Based on these features, proteins were assigned to LPXTG-anchored proteins, non-covalently cell wall–associated proteins, or lipoproteins. Proteins shown in bold were detected in at least two experimental conditions.

In the extracellular fraction, very few proteins carrying classical surface anchoring motifs were detected. No LPXTG-anchored proteins were found to be altered in the *ΔrmlT* mutant, whereas only one protein was identified as increased in abundance in the *Δlmo1079* mutant relative to the WT strain. Among proteins predicted to associate non-covalently with the cell wall, the autolysin Ami was detected as increased in abundance in both mutants compared with the WT. No lipoproteins were detected among the differentially abundant extracellular proteins in either mutant, suggesting that WTA glycosylation loss has a limited impact on proteins strongly retained at the cell surface in this fraction.

A greater number of surface-associated proteins were detected in the surface-accessible proteome. In this fraction, five LPXTG-anchored proteins were found to be increased in abundance, while two were decreased in the *ΔrmlT* mutant. Regarding *Δlmo1079* mutant, four proteins were increased and three decreased in abundance, indicating that LPXTG-anchored proteins are only moderately affected by WTA glycosylation loss. Several non-covalently associated proteins were also detected in this fraction. In the *ΔrmlT* mutant, two such proteins were increased in abundance relative to the WT, including Ami, which was consistently detected as increased across multiple fractions. In contrast, one non-covalently associated protein corresponding to the virulence factor InlB was detected at higher abundance in the WT strain than in the mutant. In *Δlmo1079*, two non-covalently associated proteins were found to be increased relative to the WT, again including Ami, suggesting that loss of WTA glycosylation enhances the accessibility of proteins relying on non-covalent interactions with the cell wall. Only two lipoproteins were detected among the differentially abundant proteins in the *Δlmo1079* mutant, with one displaying increased abundance in the mutant and the other in the WT strain. No lipoproteins were identified among the differentially abundant proteins in the *ΔrmlT* mutant, indicating that this class is only marginally affected at the level of surface accessibility.

The most pronounced differences were observed in the surface-exposed proteome. In this fraction, three LPXTG-anchored proteins were found to be altered in the *ΔrmlT* mutant, with two increased in the mutant and one more abundant in the WT strain. In *Δlmo1079*, twice as many LPXTG-anchored proteins were detected compared to ΔrmlT, including five increased in the mutant and only one increased in the WT, further supporting a limited but detectable effect on this class of proteins. Non-covalently associated proteins were also detected in this fraction, with four proteins increased in abundance in *ΔrmlT* compared with the WT, again including Ami. In *Δlmo1079*, six non-covalently associated proteins were detected, all displaying higher abundance in the mutant strain, consistent with an increased surface exposure of proteins weakly associated with the cell wall.

Strikingly, the largest group of altered proteins in the surface-exposed proteome corresponded to lipoproteins, indicating that this class is particularly sensitive to WTA glycosylation loss. In the Δ*rmlT* mutant, nine lipoproteins were detected as increased in abundance relative to the WT strain. An even stronger effect was observed in the *Δlmo1079* mutant, where 33 lipoproteins were identified as differentially abundant in the surface-exposed proteome. Among these, only one protein displayed higher abundance in the WT strain, whereas the remaining lipoproteins were consistently more abundant in the mutant, suggesting a pronounced increase in their surface exposure.

Taken together, these results indicate that the loss of WTA glycosylation differentially impacts proteins depending on their anchoring mechanisms. While only limited changes were observed for LPXTG-anchored proteins, a substantial enrichment of lipoproteins was detected among the surface-exposed proteins, particularly in the *Δlmo1079* mutant, suggesting that WTA glycosylation preferentially influences the surface exposure of lipoproteins rather than broadly affecting all classes of surface-anchored proteins.

Notably, proteins that were differentially abundant in both mutants consistently displayed the same trend, being either increased or decreased in both ΔrmlT and Δlmo1079 strains. This consistent directionality suggests that the loss of WTA glycosylation induces common structural alterations in the cell wall, likely affecting its organization or architecture, which in turn modulates protein accessibility and surface exposure.

### WTA glycosylation modulates the surface exposure of virulence-associated proteins

To determine whether the proteins altered upon WTA glycosylation loss included virulence-associated factors, the differentially abundant proteins identified in each fraction were screened for known or predicted virulence functions based on literature annotation and functional databases (**Figure 5)**.

**Figure 5.**
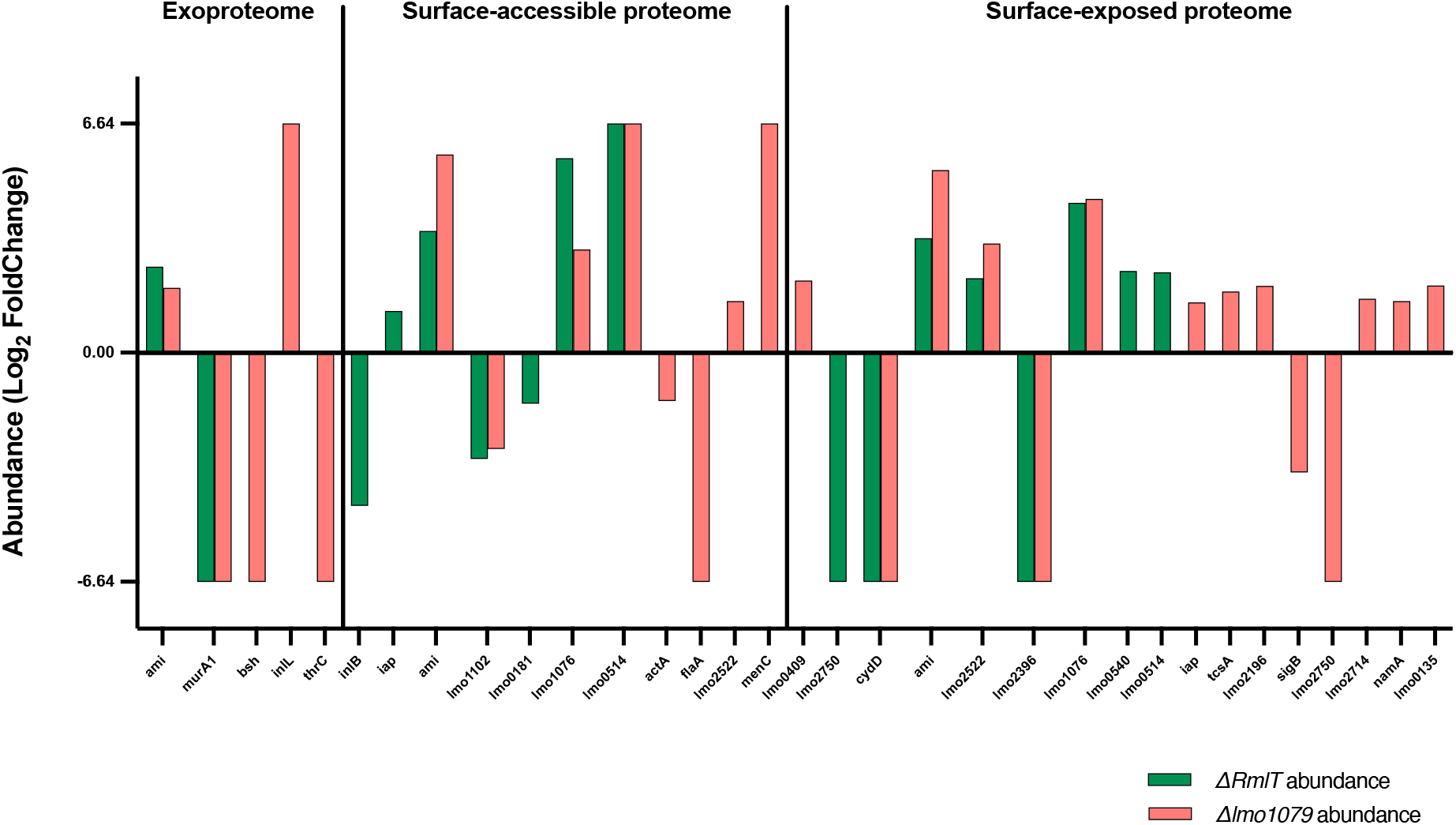
WTA glycosylation loss differentially affects virulence-associated proteins across extracellular and surface-associated fractions. Relative protein abundance of virulence-associated proteins in the extracellular, surface-accessible, and surface-exposed fractions of *ΔrmlT* and *Δlmo1079* strains vs WT strain. Positive values indicate increased abundance, whereas negative values indicate decreased abundance relative to WT. Darker shades represent *ΔrmlT*, whereas lighter shades represent *Δlmo1079*.

This allowed the distribution of virulence-associated proteins across the extracellular, surface-accessible, and surface-exposed proteomes to be assessed (**Table 2**). In the extracellular fraction, only a limited number of virulence-associated proteins were detected as differentially abundant. Among these, the autolysin Ami was consistently increased in abundance in both *ΔrmlT* and *Δlmo1079* mutants relative to the WT strain. In contrast, the cell wall–associated enzyme MurA1 was decreased in both mutants, indicating a shared response affecting proteins involved in cell envelope homeostasis. In addition, several virulence-associated proteins displayed mutant-specific patterns. In the *Δlmo1079* mutant, ThrC and Bsh were decreased in abundance, whereas the internalin-like protein InlL was increased, suggesting that individual glycosyltransferases can differentially affect specific subsets of extracellular virulence-associated proteins. Overall, these results indicate that WTA glycosylation loss has a relatively modest impact on the extracellular pool of virulence factors.

**Table 2.** Virulence-associated proteins across the extracellular, surface-accessible, and surface-exposed proteomes. Quantitative changes in protein abundance across ΔrmlT and Δlmo1079 strains. Proteins were identified in the exoproteome, surface-accessible proteome, or surface-exposed proteome, with log_2_ fold-changes indicated for each condition. Only proteins with measurable changes are reported; a dash (–) indicates the protein was not detected in that condition. Proteins shown in bold were detected in at least two conditions.

| Condition | Accession Number | Gene name | ΔRmlT | Δlmo1079 |
| --- | --- | --- | --- | --- |
| Exoproteome | **Q8Y496** | **ami** | 2.49 | 1.88 |
|  | Q8Y4C4 | *murA1* | -6.64 | -6.64 |
|  | Q8Y4A5 | *thrC* | -6.64 | - |
|  | Q8Y5J3 | *bsh* | -6.64 | - |
|  | Q8Y5N1 | *inlL* | 6.64 | - |
| Surface-accessible<br>Proteome | P0DQD2 | *inlB* | -4.44 | - |
|  | **P21171** | **iap** | 1.21 | - |
|  | **Q8Y496** | **ami** | 3.53 | 5.74 |
|  | Q8Y817 | *lmo1102* | -3.08 | -2.79 |
|  | Q8Y837 | *lmo0181* | -1.48 | - |
|  | **Q8Y842** | **lmo1076** | 5.64 | 2.99 |
|  | **Q8Y9L3** | **lmo0514** | 6.64 | 6.64 |
|  | P33379 | *ActA* | - | -1.4 |
|  | Q02551 | *flaA* | - | -6.64 |
|  | **Q8Y4C8** | **lmo2522** | - | 1.5 |
|  | Q8Y4D0 | *menC* | - | 6.64 |
|  | Q8Y9W0 | *lmo0409* | - | 2.09 |
| Surface-exposed<br>Proteome | Q8Y3T1 | *lmo2750* | -6.64 | -6.64 |
|  | Q8Y3W4 | *cydD* | -6.64 | -6.64 |
|  | **Q8Y496** | **ami** | 3.32 | 5.29 |
|  | **Q8Y4C8** | **lmo2522** | 2.16 | 3.16 |
|  | Q8Y4N9 | *lmo2396* | -6.64 | -6.64 |
|  | **Q8Y842** | **lmo1076** | 4.34 | 4.46 |
|  | Q8Y9I8 | *lmo0540* | 2.37 | - |
|  | **Q8Y9L3** | **lmo0514** | 2.33 | - |
|  | **P21171** | **iap** | - | 1.46 |
|  | Q48754 | *tcsA* | - | 1.77 |
|  | Q7AP52 | *lmo2196* | - | 1.94 |
|  | Q7AP79 | *sigB* | - | -3.47 |
|  | Q8Y3W5 | *lmo2714* | - | 1.56 |
|  | Q8Y3Y8 | *namA* | - | 1.5 |
|  | Q8Y4N9 | *lmo2396* | - | - |
|  | Q8YAJ0 | *lmo0135* | - | 1.95 |

A markedly different pattern was observed in the surface-accessible proteome. In this fraction, several well-characterized virulence-associated proteins were differentially abundant. The internalin InlB was reduced in abundance in the *ΔrmlT* mutant, whereas the autolysin Iap was slightly increased, indicating that WTA glycosylation loss can differentially affect the accessibility of key surface proteins. Notably, Ami was consistently increased in both mutants, together with *lmo1076* and *lmo0514*, both associated with the cell envelope. In particular, Ami and lmo1076 are predicted to participate in peptidoglycan remodelling, while lmo0514 encodes a surface-anchored internalin-like protein. This coordinated increase reinforces the observation that cell envelope-associated proteins are strongly affected by WTA glycosylation status. In contrast, ActA and FlaA were decreased in the *Δlmo1079* mutant, highlighting a reduction in specific virulence-associated proteins involved in actin-based motility and motility-related functions. Several additional proteins, including lmo2522 and MenC, displayed increased abundance specifically in the *Δlmo1079* mutant. Together, these results indicate that loss of WTA glycosylation alters the accessibility of virulence-associated proteins at the bacterial surface in a protein- and mutant-dependent manner.

The most pronounced changes were observed in the surface-exposed proteome. In this fraction, a strong enrichment of virulence-associated proteins was detected, particularly in the *Δlmo1079* mutant. The autolysin Ami was again consistently increased in abundance in both mutants, confirming a robust and reproducible effect across all fractions. Similarly, the proteins lmo2522 and lmo1076 were increased in both mutants, indicating enhanced surface exposure. In contrast, several proteins such as CydD and lmo2396 were consistently reduced in both mutants, suggesting decreased exposure or retention at the bacterial surface. Additional virulence-associated proteins displayed mutant-specific changes. For example, Iap was increased in the *Δlmo1079* mutant, whereas lmo1076 was strongly enriched in both mutants. Overall, the *Δlmo1079* mutant exhibited a broader and more pronounced increase in surface-exposed virulence-associated proteins compared with *ΔrmlT*.

Taken together, these results demonstrate that loss of WTA glycosylation leads to a progressive redistribution of virulence-associated proteins from the extracellular fraction towards the bacterial surface, with the most substantial effects observed in the surface-exposed proteome. While some proteins are consistently affected across both mutants, others display mutant-specific patterns, indicating that different glycosyltransferases contribute in distinct ways to the control of virulence factor accessibility and exposure at the bacterial surface. Notably, virulence-associated proteins that were altered in both mutants consistently followed the same trend, being either increased or decreased in abundance in both *ΔrmlT* and *Δlmo1079* strains. This coordinated behaviour further supports the notion that WTA glycosylation influences the overall organization of the cell wall, thereby modulating the accessibility and surface display of virulence factors

## Discussion

Our results are in line with the hypothesis that WTA glycosylation contributes broadly to the organization of *L. monocytogenes* cell envelope rather than affecting only a small subset of surface proteins. WTAs are major peptidoglycan-linked glycopolymers that extend through the Gram-positive cell wall and are known to influence cell-wall architecture, pathogenesis, and interactions with antimicrobials^6^. In *L. monocytogenes*, previous work has already shown that WTA glycosylation affects virulence and surface protein retention, particularly for non-covalently associated factors such as Ami and InlB^15^.

In this context, our data do not point to a broad disruption of the surface proteome, but rather to a redistribution of proteins across the extracellular, surface-accessible, and surface-exposed fractions following the loss of WTA glycosylation. This interpretation fits well with prior evidence that WTA decorations alter the physicochemical properties of the cell envelope. In the case of rhamnosylation, loss of RmlT increases cell-wall permeability and facilitates penetration of host antimicrobial peptides^9^ while in the case of GlcNAc loss, deletion of *lmo1079* makes the bacterial surface more hydrophilic and reduces binding of free fatty acids^21^. Together, these observations argue that WTA glycosylation modulates how molecules interact with the cell wall matrix.

In line with this global redistribution, Gene Ontology analysis revealed that the affected proteins were consistently associated with similar biological processes across all fractions and both mutants. In particular, proteins linked to primary metabolism, biosynthetic processes, and macromolecule metabolism were recurrently represented, together with molecular functions related to ion binding. Although no GO term reached statistical significance after correction, this consistent functional profile suggests that WTA glycosylation loss does not selectively impact specific pathways, but rather broadly influences the distribution of abundant and multifunctional proteins across the cell envelope. This observation is consistent with previous reports describing the presence of metabolic enzymes and other moonlighting proteins at the bacterial surface and in extracellular fractions, where their localization is often influenced by cell envelope properties rather than dedicated secretion mechanisms^22,23^.

The protein anchoring analysis suggests that these effects are not uniform across protein classes. LPXTG proteins were only modestly affected, whereas non-covalently associated proteins and, most strikingly, lipoproteins were strongly enriched in the surface-exposed fraction, especially in the Δ*lmo1079* mutant. One plausible explanation is that WTA glycosylation helps define the properties of the wall, thereby influencing which proteins remain embedded, which become more exposed, and which are more easily released. This idea is consistent with the broader literature showing that cell-envelope glycopolymers can act as structural organizers of the bacterial surface and can influence the localization or accessibility of surface-associated proteins^24^.

The stronger phenotype observed in the *Δlmo1079* mutant is particularly noteworthy. Across all fractions, this mutant consistently displayed a broader and more pronounced redistribution of proteins, especially within the surface-exposed proteome. This suggests that the loss of GlcNAc glycosylation has a greater impact on protein accessibility at the bacterial surface compared to rhamnose loss. Although the underlying mechanism cannot be directly resolved, these results support the idea that different WTA glycosylations contribute in distinct ways to the organization of the cell envelope, with GlcNAc playing a more prominent role in controlling protein exposure at the outermost surface.

The virulence-factor analysis reinforces this interpretation. Rather than simply changing total abundance, WTA glycosylation loss appears to alter the distribution and accessibilit**y** of virulence-associated proteins across fractions, with the clearest effects in the surface-exposed proteome. This is important because, for *L*.*monocytogenes*, biological activity depends not only on protein production but also on correct positioning at the host-bacterium interface. Surface localization is especially critical for adhesins, invasion factors, autolysins, and proteins that remodel the envelope during infection. Previous studies have shown that WTA glycosylation promotes proper surface association of major virulence factors, including Ami and InlB, and that disruption of specific WTA sugar decorations can impair virulence-related functions^10,25,26^.

Ami provides a particularly illustrative example in our dataset, as it was consistently increased not only in the extracellular fraction but also in the surface-accessible and surface-exposed proteomes. Rather than representing a contradictory observation, this pattern likely reflects changes in its interaction with the cell wall. Ami is a non-covalently associated autolysin carrying GW modules, and its retention at the bacterial surface is known to depend on WTA glycosylation^10^. One possible explanation is that loss of WTA glycosylation alters the cell wall in a way that both weakens Ami retention and increases its surface accessibility. As a result, the protein may become more exposed and therefore more readily detected by biotinylation and shaving approaches, while another fraction may be more easily released into the extracellular environment due to reduced binding stability.

The same logic may help explain why some virulence-associated proteins decrease in one fraction while increasing in another. For proteins whose localization depends on weak or multivalent interactions with wall polymers, altered WTA composition could shift the equilibrium between retention, exposure, and release.

The interpretation of all three datasets must take into account the known limitations of surface-proteomics approaches. Both shaving and biotinylation are powerful and complementary, but neither is perfectly selective. Importantly, the combined use of both methods provided complementary and mutually supportive information. A careful comparison of the datasets revealed that some proteins were uniquely identified by one method, reflecting differences in accessibility, detection sensitivity, or extraction principles. At the same time, a subset of proteins was consistently detected by both approaches, supporting the robustness of their identification. Notably, proteins identified in common between biotinylation and shaving datasets consistently displayed the same trend in abundance, being either increased or decreased in both conditions. This concordance strengthens confidence in the observed changes and supports the notion that WTA glycosylation loss induces coherent alterations in protein surface exposure. Cytoplasmic proteins are commonly detected in bacterial surface proteomes, either because of low-level lysis during sample handling or because some classically intracellular proteins moonlight at the cell surface or in the extracellular space^22^. In *Listeria* specifically, comparative work has shown that surface extracts are often contaminated by cytoplasmic proteins, although biotinylation generally yields lower contamination than other methods and remains complementary to shaving^27^.

The presence of cytoplasmic proteins across all datasets likely reflects a combination of minor technical carryover and the detection of moonlighting proteins. However, as similar distributions were observed across WT and mutant strains, this does not affect the overall interpretation of the results.

Overall, our results suggest that WTA glycosylation plays an important role in shaping the surface architecture of *L. monocytogenes*. Rather than simply affecting protein abundance, it appears to influence how proteins are retained, exposed, or released at the bacterial surface, likely through changes in cell-wall properties.

In this context, the loss of WTA glycosylation does not lead to a simple loss of virulence factors, but instead alters how different classes of proteins are distributed across the cell envelope. This is consistent with the limited impact observed for LPXTG proteins, the strong enrichment of lipoproteins in the surface-exposed fraction, and the recurrent redistribution of proteins such as Ami across extracellular and surface-associated fractions.

Together, all these changes are prone to influence how the bacterium interacts with its environment, including host recognition and infection processes^2^.

## Materials and Methods

### Bacterial Growth Conditions

The bacterial strains used in the current study were *Listeria monocytogenes* EGD-e (Lm), Lm *ΔrmlT* (*lmo1080* deletion mutant) and Lm *Δlmo1079* (*lmo1079* deletion mutant). Bacteria were initially grown in 5 mL BHI medium as a preculture at 37 °C with agitation (200 rpm). The preculture was then diluted 1:100 into fresh BHI and grown to late exponential phase (OD_600nm_∼0.7) under the same conditions.

### Isolation of extracellular proteins

The recovery of extracellular proteins was adapted from a previously described protocol^28^. Late exponential-phase bacteria were separated from the culture supernatant by centrifugation at 4000 g for 15 min at 4 ºC. The culture supernatant, corresponding to the same number of bacterial cells, was carefully collected and filtered through a 0.22 μm pore-size filter under vacuum to remove residual cells. The filtrate was concentrated with a polyethersulfone (PES) membrane with a 3 kDa molecular weight cut-off. To inhibit protease activity, 0.2 mM phenylmethylsulfonyl fluoride (PMSF) was added to the ultrafiltrate, followed by the addition of sodium deoxycholate (0.2 mg/mL) to support protein precipitation with trichloroacetic acid (TCA, 10% w/v) at 4 ºC overnight. After centrifugation at 20000 g for 30 min at 4 ºC, the precipitate was washed three times with ice-cold acetone and air-dried; the pellet was not brought to complete dryness to avoid difficulties in resuspending the proteins. The pellet was then solubilized in lysis buffer [50 mM Tris-HCl (pH 7.4), 100 mM NaCl] and 8 M urea was added, followed by vortexing until the pellet was fully resuspended. The solution was transferred to a 3.5 kDa MWCO dialysis membrane for protein dialysis. Dialysis was performed overnight at 4 ºC with agitation. The protein solution was further concentrated using Vivaspin 6 concentrators with a 3 kDa MWCO PES membrane at 4000 g and 4 ºC. Protein concentration was determined using the bicinchoninic acid (BCA) assay (Pierce), with bovine serum albumin (BSA) as the standard. The recovered proteins were subsequently prepared for LC-MS/MS analysis. Each experiment was performed in triplicate.

### Isolation of bacterial cell-surface proteins

The bacterial cell-surface proteome was investigated using two complementary approaches previously described in detail^28^: biotin labelling (surface-accessible proteome) and trypsin shaving (surface-exposed proteome). Briefly, for biotin labelling, late exponential-phase bacteria were harvested by centrifugation at 4000 g for 10 min at RT, then washed 3 times with PMSF Buffer (1 mM PMSF in 0.01 M PBS pH 8). After centrifugation under the same conditions, the cell pellets were weighed to ensure a consistent wet-cell mass for labelling. Surface-exposed proteins were labelled with biotin [8 mM sulfo-NHS-SS-biotin in 0.01 M PBS (pH 8)], followed by incubation for 15 min at room temperature under gentle agitation. Excess biotin was quenched with three washes using glycine buffer [0.01 M PBS pH 8, 500 mM glycine], followed by centrifugation at 4000 g during 5 m discarding the supernatant at each time. Biotinylated cells were then resuspended in Triton X-100 lysis buffer [1% (v/v) Triton X-100, 0.01 mM PBS pH 8, 1 mM PMSF] and lysed using a FastPrep-24 classic (MP Biomedicals) with two cycles of 45 s at 6.5 m/s. Cell debris was removed by centrifugation at 20000 g for 30 min at 4 ºC. Biotinylated proteins were purified by affinity chromatography in a monomeric neutravidin agarose resin (Thermo Scientific). After washing with NP-40 buffer [1% (v/v) NP-40, 0.01 mM PBS pH 8], labelled proteins were eluted with reducing buffer [62.5 mM Tris-HCl pH 6.8, 2% SDS, 20% glycerol and 50 mM dithiothreitol (DTT)] for 15 min at room temperature. This step was repeated twice, and the eluates were combined. The recovered proteins were subsequently prepared for LC-MS/MS analysis.

For trypsin shaving, bacterial cells were harvested by low-speed centrifugation at 1000 g for 15 min at 4 ºC to minimize cell lysis and washed twice with ice-cold washing buffer [20 mM Tris-HCl pH 7.4 and 150 mM NaCl]. The washed cells were resuspended in shaving buffer [20 mM Tris-HCl pH 7.4, 150 mM NaCl, 10 mM CaCl_2_ and 1 M L-arabinose] supplemented with trypsin (0.5 μg/mL). Proteolysis was performed at 37 ºC for 1h with gentle shaking to cleave surface accessible proteins. After digestion, cells were pelleted by centrifugation (1000 g, 15 min, 4 ºC), and the supernatant containing cleaved peptides was filtered through a 0.22 μm pore-size filter to remove bacterial debris. The resulting peptide mixture was then submitted for downstream proteomic analysis by LC-MS/MS.

Each experiment was performed in triplicate for both approaches.

### Proteomic Sample Preparation and LC-MS/MS Acquisition

Each sample was processed for proteomic analysis following the solid-phase-enhanced sample-preparation (SP3) protocol and enzymatically digested with trypsin/LysC as previously described^29^. Protein identification and quantitation was performed by nanoLC-MS/MS equipped with a Field Asymmetric Ion Mobility Spectrometry - FAIMS interface. This equipment is composed of a Vanquish Neo liquid chromatography system coupled to an Eclipse Tribrid Quadrupole, Orbitrap, Ion Trap mass spectrometer (Thermo Scientific, San Jose, CA). 250 nanograms of peptides of each sample were loaded onto a trapping cartridge (PepMap Neo C18, 300 μm x 5 mm i.d., 174500, Thermo Scientific, Bremen, Germany). Next, the trap column was switched in-line to an Aurora Frontier XT 60 cm, 75μm (AUR4-60075C18-XT) chromatographic separation column. A 116 min separation was achieved by mixing A: 0.1% FA and B: 100% ACN, 0.1% FA with the following gradient at a flow of 250 nL/min: 2 min (0% B to 4% B), 20 min (4% B to 12% B), 65 min (12% B to 28% B), 11 min (28% B to 45% B), 2 min (45% B to 85 % B) and 16 min at 99% B. Subsequently, the column was equilibrated with 0% B. Data acquisition was controlled by Xcalibur 4.7 and Tune 4.2.4321 software (Thermo Scientific, Bremen, Germany). MS results were obtained following a Data Dependent Acquisition - DDA procedure. MS acquisition was performed with the Orbitrap detector at 120 000 resolution in positive mode, quadrupole isolation, scan range (m/z) 375-1500, RF Lens 30%, standard AGC target, maximum injection time was set to auto, 1 microscan, data type profile and without source fragmentation.

FAIMS mode: standard resolution, total carrier gas flow: static 4L/min, FAIMS CV: -45, -60 and -75 (cycle time, 1 s). Internal Mass calibration: Run-Start Easy-IC. Filters: MIPS, monoisotopic peak determination: peptide, charge state: 2-7, dynamic exclusion 30s, intensity threshold, 5.0×10^3^.

MS/MS data acquisition parameters: quadrupole isolation window 1.8 (m/z), activation type: HCD (30% CE), detector: ion trap, IT scan rate: rapid, mass range: normal, scan range mode: auto, normalized AGC target 100%, maximum injection time: 35 ms, data type centroid.

### Protein Identification, Quantification, and Data Analysis

The raw data was processed using the Proteome Discoverer 3.1.1.93 software (Thermo Scientific) and searched against *L. monocytogenes* 1/2a strain ATCC BAA-679/EGD-e reference proteome (taxonomy ID 169963) retrieved from the UniProt database. A common protein contaminant list from MaxQuant was also included in the analysis. The Sequest HT search engine was used to identify tryptic peptides. The ion mass tolerance was 10 ppm for precursor ions and 0.5 Da for fragment ions. The maximum allowed missing cleavage sites was set to two. Cysteine carbamidomethylation was defined as constant modification. Methionine oxidation, deamidation of glutamine and asparagine, peptide terminus glutamine to pyroglutamate, and protein N-terminus acetylation, Met-loss, and Met-loss+acetyl were defined as variable modifications. Peptide confidence was set to high. The processing node Percolator was enabled with the following settings: maximum delta Cn 0.05; target FDR (strict) was set to 0.01 and target FDR (relaxed) was set to 0.05, validation based on q-value. Protein label-free quantitation was performed with the Minora feature detector node at the processing step. Precursor ions quantification was performed at the consensus step with the following parameters: inclusion of unique plus razor peptides, precursor abundance based on intensity, and normalization based on total peptide amount. For hypothesis testing, protein ratio calculation was pairwise ratio-based and an t-test (background based) hypothesis test was performed.

### Bioinformatic Analysis of the Secretome and Surfaceome

Secretomic analysis was conducted to evaluate whether the proteins identified by mass spectrometry were plausibly extracellular or surface-associated proteins. A comprehensive bioinformatic pipeline integrating multiple prediction tools was applied to infer secretion pathways, subcellular localization, and surface anchoring. Classical secretion signals were predicted using SignalP v6.0^30^, which detects Sec, Tat, and lipoprotein signal peptides. Non-classically secreted proteins were identified with DeepLocPro v1.0^31^ (prokaryotic model), which predicts subcellular localization independently of signal peptide presence. DeepTMHMM v1.0^32^ was used to predict transmembrane helices, enabling the distinction between transmembrane and truly secreted proteins lacking membrane-spanning regions.

Surface anchoring motifs (LPXTG and other sortase-dependent signals) were predicted using GPAPred^33^. All prediction results were integrated into a unified dataset (**Table S1**) containing all proteins validated by mass spectrometry (FDR ≤ 0.01 and >2 peptides). For each protein, secretion pathway and subcellular localization predictions were annotated, allowing direct comparison between experimental detections and bioinformatic inferences. This integrated approach confirmed the presence of extracellular and surface-associated proteins, while acknowledging the expected occurrence of cytoplasmic contamination typical of these sample preparations. To account for uncertainty in subcellular localisation prediction, the complete protein dataset was screened against the MoonProt 3.0^34^ database, to identify proteins with reported moonlighting activities and multifunctional roles. Gene Ontology (GO) terms for molecular function and biological process were obtained using the DAVID^35^ functional annotation platform, applying an EASE threshold of 1 and a count threshold of 2.

## Author’s contributions

RM and DC conceptualise the overarching aims of the research study. GM, RM and DC conceived and designed the experiments. GM performed the experiments and data acquisition. GM, RM and DC analysed and interpreted the data. RM and DC had management as well as coordination responsibility for the execution of the research work. RM and DC contributed to the acquisition of the financial supports and resources leading to this publication. All authors contributed to the drafting of some parts of the manuscript, including reading and revising critically the manuscript for important intellectual content, as well as approval of the final version.

